# The First Comprehensive Description of the Platelet Single Cell Transcriptome

**DOI:** 10.1101/2024.10.15.618506

**Authors:** Walter Wolfsberger, Chase Dietz, Caitlin Foster, Taras Oleksyk, A. Valance Washington, Donald Lynch

**Affiliations:** Department of Biological Sciences, Oakland University. Rochester MI 48309; University of Cincinnati College of Medicine - Center for Cardiovascular Research

**Keywords:** Platelets, Transcriptomics, RNA, Single cell sequencing, Megakaryocytes, Thrombosis

## Abstract

Platelets are derived from megakaryocytes, either in peripheral or pulmonic circulation. The transcriptome of megakaryocytes has been studied, while the platelet transcriptome is thought to be a reflection of their parent cells; it has not yet been investigated. Although platelets lack nuclei, they inherit RNA from their parent megakaryocytes, while only about 10% of them are believed to contain enough RNA for meaningful analysis. This study explores the potential of single-cell RNA sequencing to analyze the platelet transcriptome, aiming to expand our understanding of platelets beyond their traditional role in coagulation. Using acridine orange staining and antibody-based sequencing, we successfully sequenced RNA from seven healthy donors. Results revealed significant heterogeneity in gene expression, with common platelet markers, such as *ITGA2B* and GP1B, being less abundant than expected. Interestingly, immune markers associated with lung megakaryocytes were not strongly represented in peripheral platelets. Comparison with current algorithms for cell identification suggests that platelets are often misclassified as other blood cell types, highlighting limitations of existing pipelines in platelet annotation. This misclassification may have led to misrepresentation of platelet transcriptomics in previous studies. These findings underscore the need for tailored sequencing methods to accurately profile platelets and set the foundation for further exploration of platelet biology and immune function, potentially opening avenues for therapeutic interventions in immune modulation, drug delivery, and the use of platelets as disease biomarkers in cancer and other conditions.

**Key Points:** Platelet single cell sequencing can be implemented with appropriate technical refinements to ensure optimal isolation without exogenous activation. In comparison to bulk sequencing techniques, single cell analysis affords the ability to exclude contaminating cells enabling examination of the authentic platelet transcriptome. This is critically important as contaminating cells contain far more RNA ultimately skewing results of transcriptomic analysis.

Most platelets do not contain significant levels of commonly expected transcripts such as ITGA2B, GP1B, TREML1. In the context of recent data, our single cell transcriptomic data supports the intradividual and interindividual heterogeneity of the platelet transcriptome. The lung megakaryocyte signature is not disguisable in peripheral platelets. Further studies are needed to understand sources of RNA within platelets and the impact of the platelet microenvironment.

Visual abstract

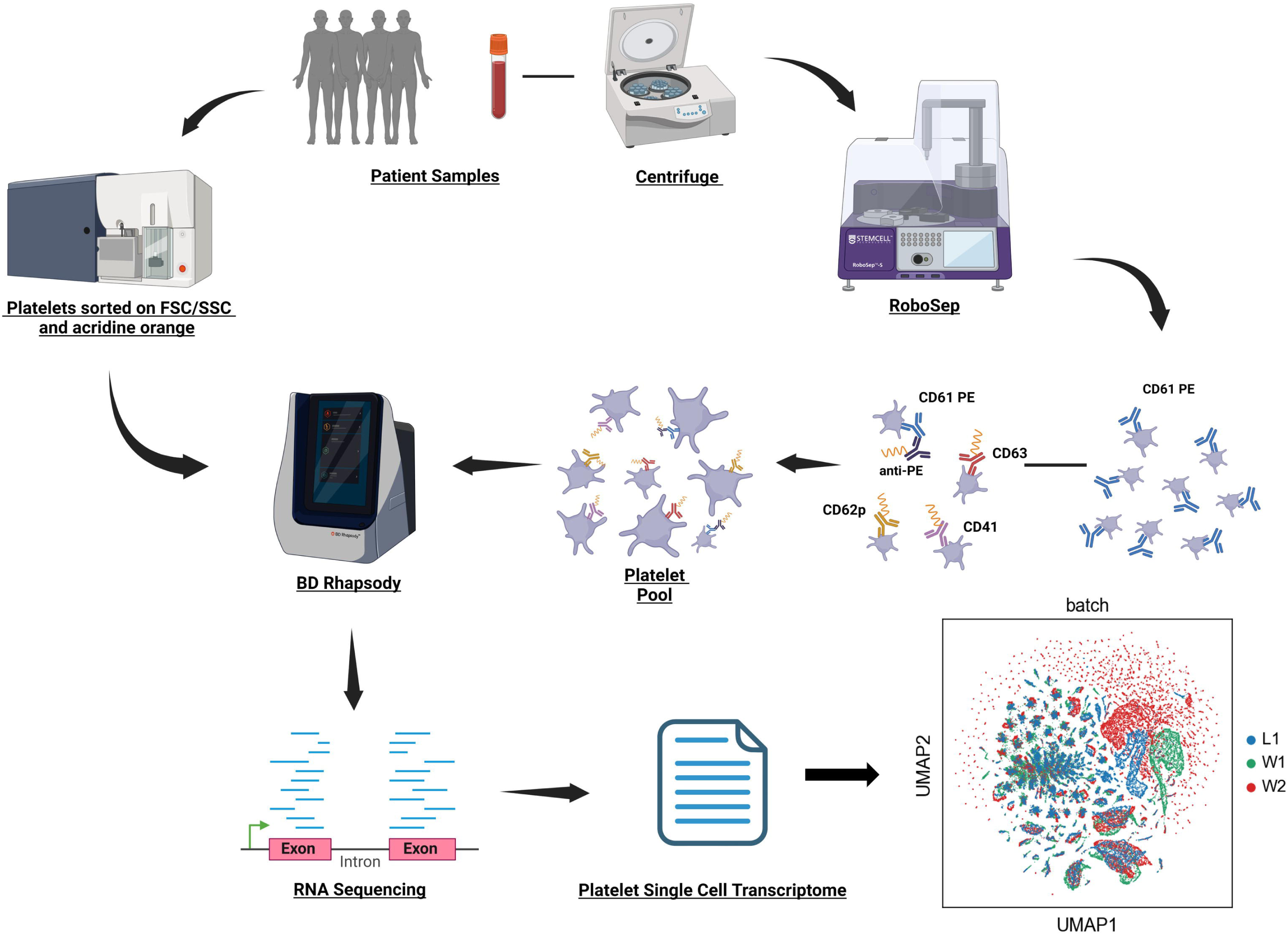

## Introduction

Platelets are small, anuclear cells derived from megakaryocytes and are the second most abundant cell type in peripheral blood. Since their discovery in the late 1800s, platelets have been primarily associated with maintaining vascular integrity through the formation of blood clots. However, recent research has uncovered a broader role for platelets in immune function. For example, platelets possess toll-like receptors that detect pathogen-associated molecular patterns (PAMPs), such as LPS, CpG islands, DNA, and RNA. Additionally, platelets store and release cytokines that modulate the immune responses(1, 2).

Although platelets lack a nucleus, they inherit RNA from their parent megakaryocytes and have the ability to translate this RNA into proteins. Interestingly, studies have shown that megakaryocytes, particularly in the lungs, express major histocompatibility complex (MHC) class II, a feature previously thought to be restricted to immune-regulating cells like macrophages, dendritic cells, and lymphocytes. Recent evidence also suggests that a population of bone marrow megakaryocytes express MHC class II, although platelets themselves are thought not to(3, 4).

Sequencing studies of megakaryocytes in humans and mouse models have revealed that these cells change their gene expression profiles in response to disease. For instance, during infections with viruses like dengue and influenza, several megakaryocyte transcripts, including *IFITM3*, are upregulated, correlating with better survival outcomes(5). However, it remains unclear whether these transcriptional changes are passed on to platelets and whether they influence platelet function. Exploring the platelet transcriptome could provide new insights into these questions, though the first step is analyzing platelet RNA at the single-cell level.

We propose using single-cell RNA sequencing to investigate the RNA platelets inherit from megakaryocytes, which may reveal their potential as small reservoirs of immune-related molecules. This knowledge could enable the manipulation of platelets for therapeutic interventions, including immune modulation and drug delivery. While we have some understanding of the proteins platelets carry, their transcriptome remains poorly characterized, representing a significant gap in our knowledge of platelet biology and immune function.

A major challenge is that only around 10% of platelets are believed to contain RNA, with those that do, carrying approximately 2.2 femtomoles of RNA per platelet(6), making sequencing difficult and risky. To date, platelet sequencing has been limited to bulk RNA sequencing or by the identification of platelets through the presence of surface markers such as CD41. These methods assume that the transcriptome contains enough information to accurately identify platelets within mixed cell populations.

In our study, we applied single-cell RNA sequencing and antibody-based sequencing (AbSeq) to successfully perform platelet transcriptomic analysis and found significant heterogeneity in RNA expression between platelets. Surprisingly, transcripts for common platelet proteins were much scarcer than expected. Furthermore, the unique immune signature of lung megakaryocytes reported in the literature was not easily detectable in peripheral platelets of humans, highlighting the gaps in our current understanding. We found that the majority of the platelets were misidentified as other cell types using current pipelines for cell annotation.

## Materials and Methods

### Platelet isolation

All human subject samples were collected under the regulations of the declaration of Helsinki and evaluated by the Institutional Review Board at Oakland University or the University of Cincinnati. All volunteers provided informed consent prior to blood collection (IRB-FY2021-390 - Translational studies on the platelet receptor TLT-1). All authors had access to the primary clinical data and it was analyzed by WW and DL. For the acridine orange experiments, platelets were collected in sodium citrate tubes and isolated from individuals after informed consent using previously published protocols(7). For acridine orange studies, the three donors were: a healthy Caucasian female aged 27, a healthy Caucasian female aged 47, and a healthy Afro-American male aged 58 each Platelets were stained with acridine orange (AO) by mixing platelets at 2 x 10^5^/ml 4:1 with freshly made AO staining buffer (AO 25μg/ml, 0.1M citric acid, 200 mM Na phosphate) for ½ hour at room temperature and sorted directly on a FACS Aria II based on the forward and side scatter of platelets and acridine orange high. For the AbSeq experiments: whole blood samples were obtained from four healthy donors and processed within 30 minutes following collection. Donors consisted of two healthy females and two healthy males, ages 21 – 55, with each being from unique ethnic groups. Platelet-rich plasma was isolated following centrifugation at 200g for 17 minutes. Platelets were subsequently isolated following centrifugation at 1,200g for 8 minutes after which the pellet was resuspended in 3 mL of PIPES/saline/glucose (PSG) buffer. Platelets were isolated using the Robosep™-S (Stemcell Technologies, Vancouver, Canada; catalog no. 21000) with use of EasySep™ Human CD45 Depletion Cocktail (Stemcell Technologies; catalog no. 17898C) and EasySep™ RBC Depletion Reagent for Robosep™ (Stemcell Technologies; catalog no. 18170) reagent kits for negative isolation of untouched platelets. For clarity, figure 1 is a schematic of the acridine orange and AbSeq designs. The purity of isolated platelets was evaluated with flow cytometry using CytoFlex (Beckman Coulter, USA).

**Figure 1.**
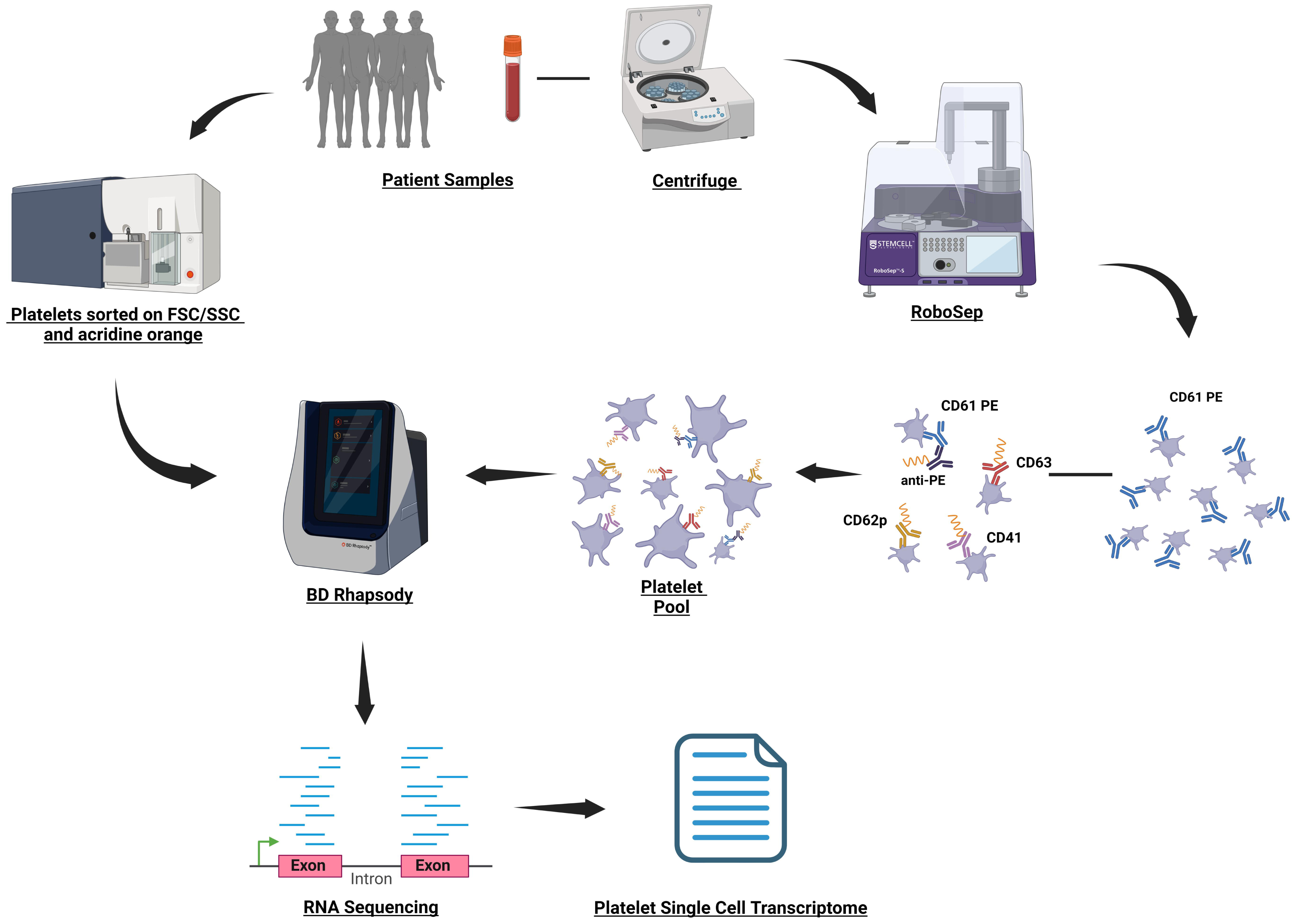
– Platelet single cell RNA sequencing experimental design: Schematic showing the strategies used for platelet single cell sequencing using either: enrichment of acridine orange positive cells with cell sorting or surface labeling with Abseq oligos targeting platelet surface markers. BD Rhapsody was used for cell capture which was followed by library prep and sequencing.

### cDNA Library production

cDNA was made according to the Rhapsody™ System Single-Cell Labeling with BD® Single-Cell Multiplexing Kit and BD® AbSeq Ab-Oligos protocol. Briefly, each sample was labeled with CD61-PE (BD Biosciences; catalog no. 561912) for 30 minutes after which 2 washes were performed in Tyrode’s buffer. Each of the four samples was labeled with a unique sample tag from the Flex Single-Cell Multiplexing Kit A (BD Biosciences; catalog no. 633849). The Ab-Oligos mouse anti-human CD41a, mouse anti-human CD62P, and mouse anti-human CD63 were used to stain samples for further proteomic labeling. Approximately 2 million platelets from each sample were combined to create a pool which was used for cell capture and subsequent steps.

### Cell Capture

Platelets were held at room temperature until cell capture. Platelets were labeled with 9.9 uM Calcein AM (BD Bioscienes, New Jersey, catalog no 564061) Platelets were captured using the Rhapsody™ HT Single-Cell Analysis System and protocol (BD Biosciences, New Jersey, USA). Imaging was performed using the Rhapsody scanner to evaluate capture efficiency, as shown in Figure 1. Capture efficiency was calculated using Rhapsody software and verified manually from the brightfield image.

### Library Preparation

Using captured platelets from four healthy donors, a library was prepared using the Rhapsody™ System mRNA Whole Transcriptome Analysis protocol with the Rhapsody™ WTA Amplification Kit (BD Biosciences; catalog no. 633801) and AMPure® XP magnetic beads (Beckman Coulter, California, USA; catalog no. A63881). QC was performed using Tapestation (D1000) along with Quibit 2. The resulting cDNA libraries were sequenced (acridine orange samples sequenced at UCLA core sequencing laboratory and AbSeq samples sequenced at the University of Cincinnati).

### Sequencing

The libraries were pooled at equimolar concentrations after which they were sequenced on a NovaSeq flow cells (Illumina, California, USA). The acridine orange samples were sequenced at UCLA core sequencing laboratory using 3 S1 lanes (Illumina, California, USA). The AbSeq samples were sequenced at the University of Cincinnati on a NextSeq P2 flow cell (Illumina, California, USA).

### Data analysis

Sequencing data was first analyzed using the Seven Bridges platform for the Rhapsody produced cDNA libraries using revision 16, version 2.2.1 of BD Rhapsody™ Sequence Analysis Pipeline using *RhapRef_Human_WTA_2023-02* reference dataset(8). The acridine orange-stained experiments were ran with Exact Cell Count set to 50000. Primary analysis output in h5ad format was analyzed using Scanpy single-cell analysis library with Python programming language(9). For every individual experiment; we filtered out cells that contained any expression of PTPRC, HBB, HBA1, HBA2, HBD, and GYPC to remove contaminating leukocyte and erythroid cells (Supplemental figure 1). We also excluded long non-coding RNA (lncRNA) genes based on the Ensembl *hsapiens* gene database. Additionally, we removed genes demonstrating sex-biased expression using a curated list (10), to minimize potential confounding effects of sex differences on downstream analyses. We calculated the qc metrics and plotted them as violin plots after filtering out genes present less than in 5 cells. Outlier cells in the data were removed based on Median Absolute Deviation (MAD) value of 3 for the log-transformed number of cells, total UMI per cell, and percentage of mitochondrial counts per cell. Due to the experiment’s focus on platelets, cells with higher than usual % of mitochondrial content were retained at <30% threshold. Total counts were normalized with the sum of 10,000 counts per cell and log transformed. Doublet detection was performed using scrublet (11). At this step, we branched our analysis – with one approach using the Seurat method to detect the top 3000 highly variable genes (12), while regressing out unwanted sources of variation, and the other, more permissive, retaining all the genes. Data was scaled to unit variance and zero mean. We computed the nearest neighbors distance matrix and a neighborhood graph of observations (13) for UMAP and cluster the cells using the Leiden algorithm with moderate resolution at different parameter values. The heterogeneity of platelets does not allow us to use high resolution clustering, we evaluated our data with different resolutions to see if we can identify novel trends. We used the Wilcoxon rank-sum (Mann-Whitney-U) and logistic regression approach (14) to rank the differentiating genes and produce plots representing the data. Decoupler-py (15) was used for cell type annotation from marker genes and functional enrichment of biological terms using the Enrichment with Over Representation Analysis. *Omnipath PanglaoDB* (*16*), a database of cell type markers was used for cell-type inference, while Molecular Signatures Database (MSigDB) GO: Gene Ontology gene sets (17, 18) was used for the functional enrichment of biological terms analysis. For data integration of multiple samples, we first merged the dataset, visualized their cell distribution using PCA to identify any significant batch effects, and later used Harmony algorithm post merge to correct for them (19). Additionally, for the integrated dataset, we used CellTypist to produce alternative cell-type annotation with *Healthy_COVID19_PBMC* reference dataset (20).

All data is available on the xxx website

## Results

### Platelet RNAseq

Since platelets lack a nucleus, it is generally believed that they are released from megakaryocytes with all the RNA they will ever contain, and this RNA content declines over the platelet’s lifespan. As a result, it is estimated that only about 10% of platelets have appreciable amounts of RNA, which has led to skepticism about the feasibility of platelet transcriptomics(6). To address the concern that most platelets may not have sufficient RNA for meaningful analysis, we used acridine orange staining to identify platelets with the highest RNA content for transcriptomic profiling.

Platelets were isolated as described in the materials and methods section, stained with acridine orange, and sorted based on scatter, followed by a single-cell gate. A distinct population of approximately 10% of acridine orange-positive platelets was separated and sorted for sequencing. In the first three experiments, varying numbers of platelets fitting the gate criteria were isolated (Figure 2A). These platelets were processed using standard BD Rhapsody™ protocols, resulting in successful sequencing from three healthy donors (HD). The sequencing results, visualized through Rhapsody™ t-distributed stochastic neighbor embedding (t-SNE) plots, showed an even distribution of transcripts across cells with some but not significant clustering. Leveraging on the ability of single cell to post-hoc remove contaminating cells, we censored out any leukocytes by identifying cells with PTPRC. Furthermore, we removed any contaminating red blood cells based on the presence of HBB, HBA1, HBA2, HBD, and GYPC leaving only platelets for further analysis (Figure 2B). Other cell types, especially those with a nucleus, would have exhibited significantly higher RNA expression levels.

**Figure 2.**
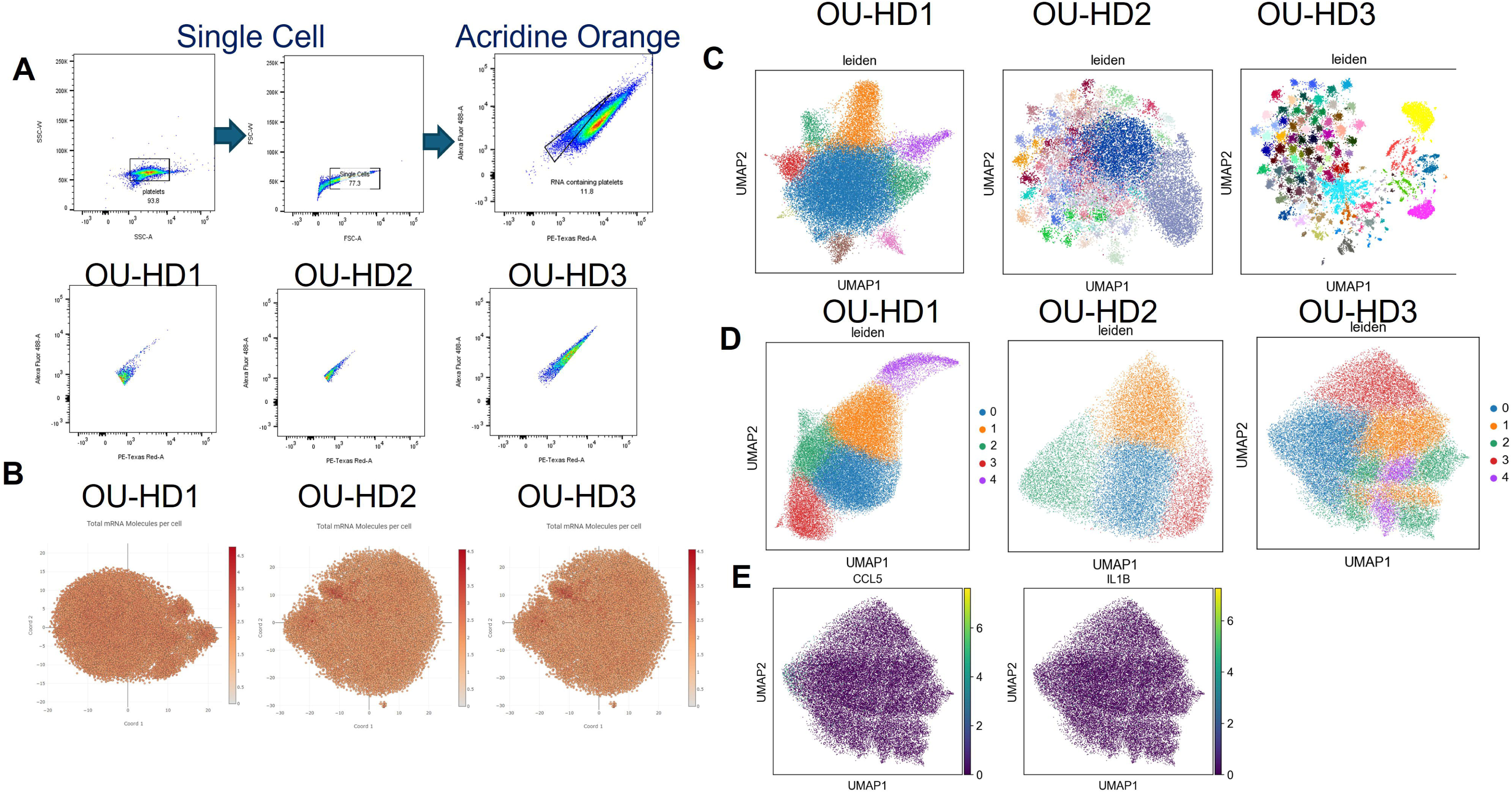
– Platelet single cell RNA sequencing: A) Gating scheme to isolate acridine orange high platelets (top) and isolated cells in the platelet gate for healthy donors 1 – 3 from left to right (bottom). B) Nearest neighbor plot showing total mRNA molecules per cell for healthy donors 1 – 3 from left to right. C, D) UMAP of each of the healthy donors 1 −3 from left to right at high stringency (C) and low stringency (D). E) Identification of immune genes which are expressed higher in lung megakaryocytes compared to bone marrow megakaryocytes.

Transcriptome analysis of 50,000 platelets yielded 2.56 billion reads, an average of 221 molecules per cell, and a median of 628 reads per cell for the first individual. The second individual generated 1.76 billion reads with an average of 266 molecules per cell, median of 5,626 reads per cell, while the third individual produced 2.4 billion reads with an average of 109 molecules per cell and a median of 7,743 reads per cell. AbSeq sample sequencing resulted in 21,911 putative cells, yielded 0.5 billion reads, with 158 molecules per cell and a median of 2,378 reads per cell. The overall quality of the reads varied, ranging from 47% in the first sample, 80% in the second and third samples, The AbSeq sampled yielded a quality of 87%. A large number of mitochondrial genes were present in all samples, which were used to QC filter the cells prior to the final analysis as described in the methods section. For the three acridine orange single-cell RNA sequencing experiments, the initial 50,000 cell count was filtered down to 34,977, 20,533 and 26,565 cells respectively with 3000 genes, while permissive filtering retained approximately 20,000 genes for each experiment. AbSeq sample filtering resulted in 16,965 cells with 3,000, or permissively 13,637 genes. UMAP representations of the data demonstrate the heterogenicity of each of the platelet samples. At a normal resolution, with filtering for variable genes, the Leiden clustering at the default resolution produced over 50 individual clusters for each individual indicating high heterogeneity of the samples. To reduce the number of clusters, we visualized the UMAP at leiden resolution parameters of0.5 and 0.3, with 0.3 results presented in Figure 2C. Due to the high number of clusters, we evaluated the results of permissive analysis, without the detection of the top 3000 variable genes, which lead to considerably less clusters in all three experiments (figure 2D). We subsequently evaluated the clusters for a lung signature based on the reports of lung megakaryocytes having a more immune profile compared to bone marrow megakaryocytes(4). We looked for transcripts that identified part of the immune profile found in megakaryocytes. However, we were unable to find any appreciable clusters that had high levels of these transcripts. We were only able to identify CCL5 and IL1B, however, they did not fit into any specific cluster that would serve as a lung phenotype (Figure 2E). To understand whether our platelet sequencing was representative of a platelet population, we compared our data to previously published bulk transcriptomic data of 204 healthy individuals (21). We found that our top transcripts matched their published results with a deviation in HD1 and samples two and three showing a marked representation as shown in Table 1. While HD1 expressed all but one of these genes, they were not as abundant as in HD 2 and 3.

**Table 1.** – Identification of the most highly ranked 14 transcripts from bulk sequencing in our single cell sequencing cohort.

| Experiment | 1 | 2 | 3 |
| --- | --- | --- | --- |
|  | Raw Molecule RANK |  |  |
| B2M | 21 | 21 | 3 |
| TMSB4X | 28 | 1 | 7 |
| FTL | 1172 | 66 | 80 |
| PF4 | 4301 | 300 | 177 |
| PTMA | 188 | 31 | 82 |
| OAZ1 | 1157 | 55 | 167 |
| PPBP | 1132 | 48 | 37 |
| ACTB | 236 | 11 | 23 |
| CLU | 1009 | 52 | 187 |
| F13A1 | 716 | 39 | 223 |
| SPARC | 515 | 34 | 69 |
| TAGLN2 | 358 | 23 | 34 |
| OST4 | NF | 24 | 19 |
| GNAS | 236 | 18 | 27 |

### Annotation of Platelet transcriptome

The cloud-based Seven Bridges sequencing report, did not consider platelets as a cell type option, classifying them as monocytes or lymphocytes. With cell-type annotation based on reference marker sets using 2 instruments, *CellTypist and* ORA analysis of *PanglaoDB markers*, only a subset of platelets was correctly labeled, while others were misidentified as B cells, dendritic cells, endothelial cells, and macrophages. The results of marker-based ORA analysis of cells filtered for top 3000 informative markers are presented in Figure 3A-C.

**Figure 3.**
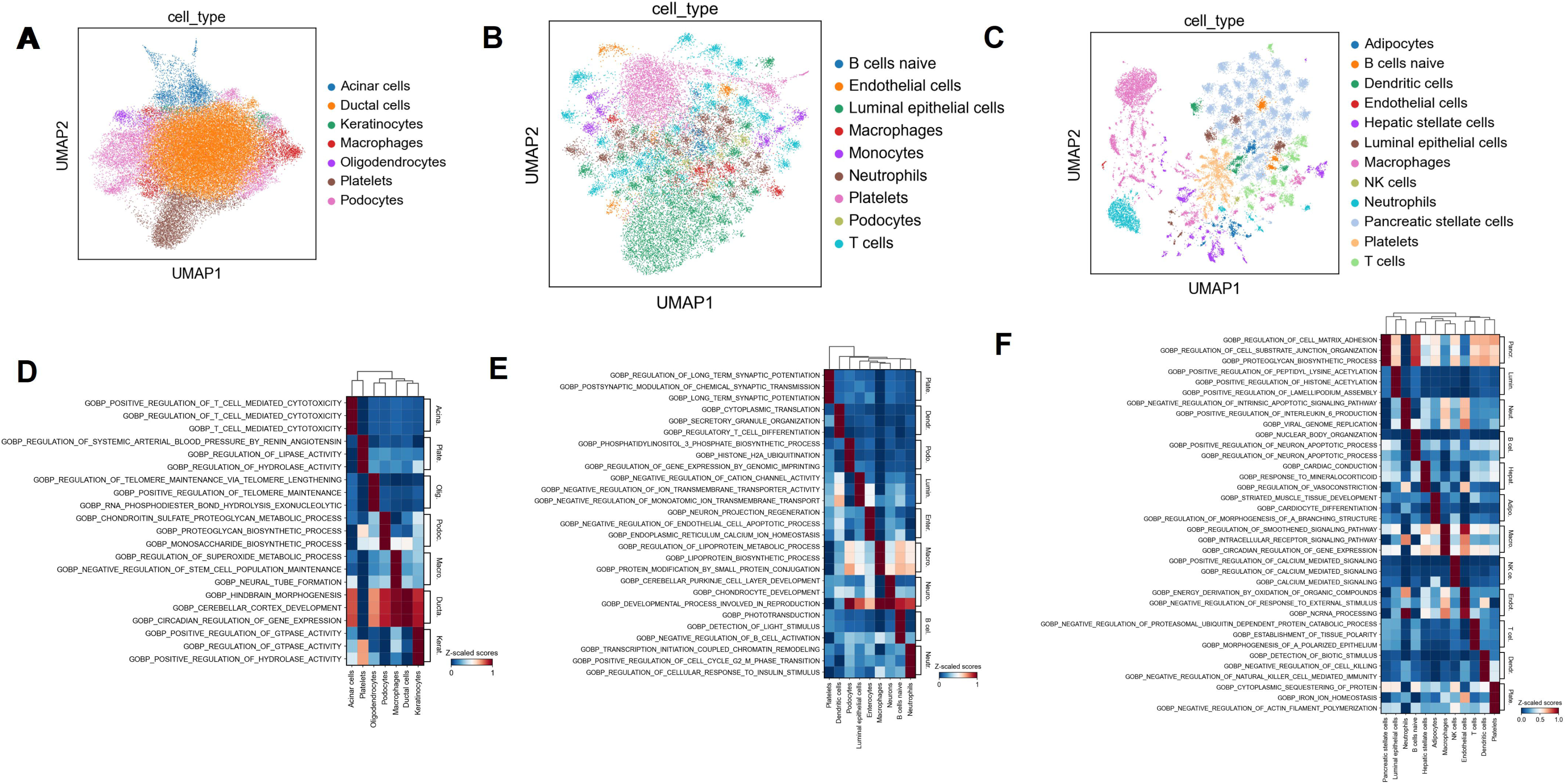
– Functional annotation analysis of the top three pathways from annotated cells and its characterization of platelets based on Gene Ontology Terms (GO). A-C) UMAP analysis of platelets based on common annotations from MSigDB. D-F) Three-dimensional heat maps showing functional annotation of platelet transcripts from HD1 (D), HD2 (E), HD3 (F), there and common cellular associations.

Functional enrichment of biological terms analysis demonstrated heterogeneity based on the clustering of the cells. The sample clusters were particularly differentiated by enrichment in pathways related to synaptic potentiation and transmission, ion hemostasis, actin filament polymerization, protein clustering, blood pressure regulation by renin angiotensin, and regulation of lipase and hydrolase activity. Other highly annotated platelet pathways included regulation of GTPase activity, cell matrix adhesion, and proteoglycan biosynthetic processes. (Figure 3D-F). Surprisingly, coagulation and immune related pathways were not among the top associated pathways in each of these samples.

### Platelet AbSeq

We were surprised to observe that key platelet transcripts, such as ITGA2B, ITGA3B, GP1B, *TREML1*, were present but exhibited low expression levels across individual platelets. To validate these findings, we employed AbSeq where cells were labeled with oligonucleotide-conjugated antibodies to ensure accurate platelet identification sequence derivation (Figure 1). Figure 4 presents the results from four healthy individuals, with platelets identified by CD41 and CD61 (*ITGA2B* and *ITGB3*) and further validated using antibodies against CD63 and CD62P as markers of activation. Through multiplexing, we obtained a total of 21,000 cells, generating 109 million reads and averaging 158 unique transcripts per cell. Specifically, the four samples yielded 8,984, 1,291, 3,873, and 3,153 cells, respectively. Rhapsody t-distributed stochastic neighbor embedding (t-SNE) analysis revealed that most cells maintained relatively consistent transcript levels, except for a small cluster in the upper right corner of Figure 4A, which exhibited lower transcript amounts. UMAP analysis highlighted heterogeneity both between individuals and within each sample. Figures 4B and 4C illustrate the numerous clusters identified in these samples; to simplify the UMAP visualization, we relaxed the parameter stringency shown in Figure 4D. Figures 4D - F demonstrate that, although these cells were identified using specific antibodies for protein targets, less than 0.2% contained CD63 and <1% contained *ITGA2B* corresponding transcripts. Additionally, SELP corresponding to CD62P was detected in less than 10 cells. These results corroborate our findings from the initial three donors, indicating that *ITGA2B* and the other genes targeted by ab-seq were minimally expressed in these samples. After mitochondrial filtering, the most highly expressed gene was the scaffolding protein *GRIP1*.

**Figure 4.**
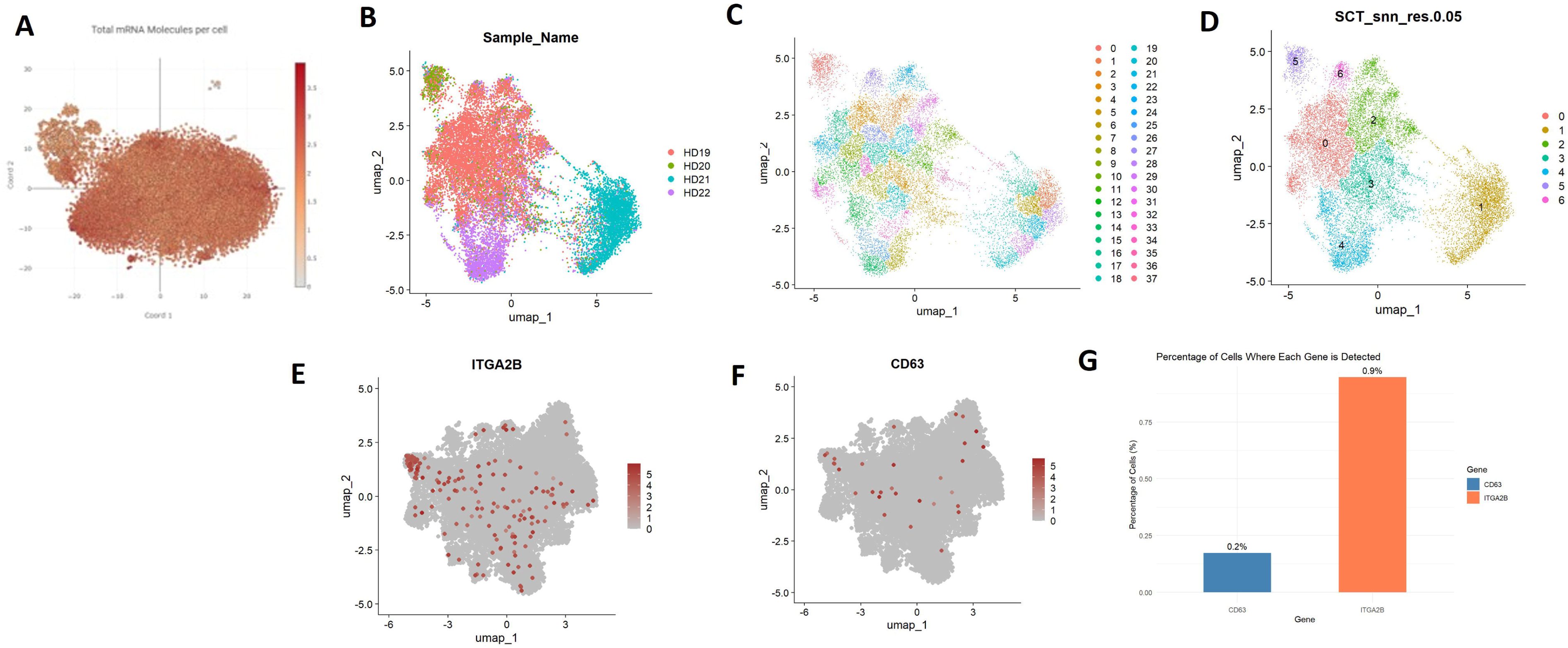
– Abseq of four healthy donors: A) Total mRNA molecules per cell for the four healthy donors evaluated by Abseq censored based on the presence of CD41 oligo, B) UMAP of the four healthy donors coded by color. C) Hi stringency UMAP (D) and low stringency UMAP of the four healthy donors. E) Correlation of CD41 protein expression with ITGA2B mRNA expression. F) Correlation of CD63 protein expression with CD63 mRNA within merged dataset. G) Bar plot of the cell level % expression of mRNA for CD63 and ITGA2B relative to protein levels.

### Experiment data integration and comparison

During our preliminary integrations, we evaluated our combined set for key platelet markers and contamination. We utilized Seurat’s scale data function Feature Plot. We applied it to the integrated data set restricting the scaling to the identified marker genes to ensure that the expression values were standardized across cells to allow accurate comparison. We identified nine distinct clusters, as shown in Supplemental figure 1A. The clusters reveal a progression from cells at the top that were not selected by acridine orange to those at the bottom that resulted in a population of acridine orange-high platelets (OU-HD 1-3: Supplemental figure 2B). In Supplemental figure 1C, we show before and after filtering of those cells that express the leukocyte contamination marker, PTPRC, along with RBC marker genes after their removal (Supplemental figure 1D). Using this population, we analyzed the highly expressed genes identified through bulk RNA sequencing from prior literature, as shown in Figure 1E. Among the most highly expressed genes were *B2M*, *TMSB4X*, and *OST4*, while *PF4* and *PPBP* were less expressed compared to the initial sample set. MHC class II genes were present, but not abundant as shown in Supplemental figure 2

For the final integration, we have selected OU-HD sample 2 and 3, and AbSeq samples due to higher values of median read per cell contributing to more reliable findings. At this step, we adopted 3 strategies for data integration. For the first approach, we merged the datasets which were pre-selected for informative variable genes (Figure 5A), for the second one we combined the results of permissive filtering of experiments and then identified the top 3000 highly variable genes within the integrated data (Figure 5B). Lastly, we analyzed the merged file of the 3 datasets with permissive filtering (Figure 5C).

**Figure 5.**
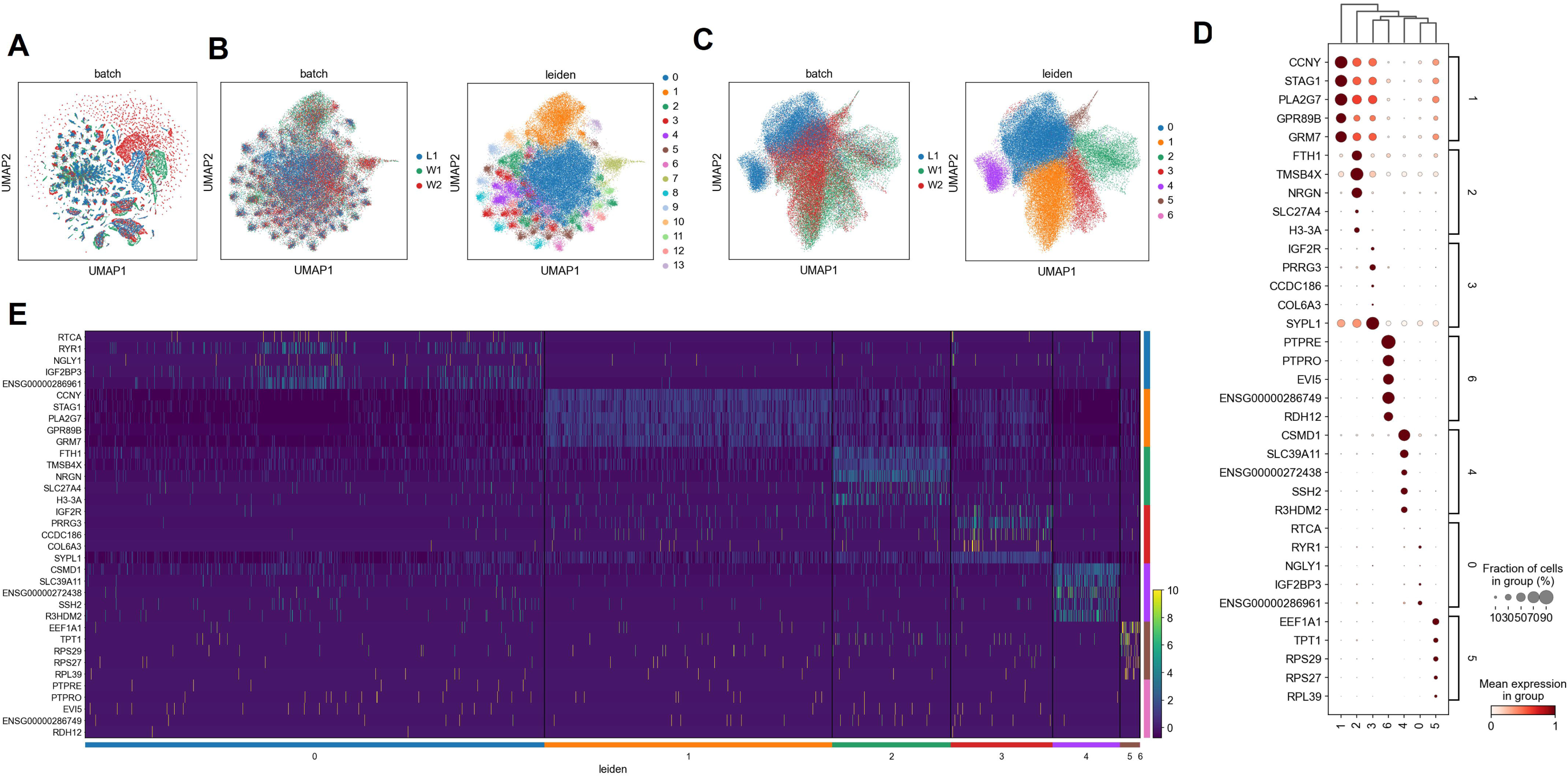
– Combined scSeq and Ab-scSeq of Platelets from Healthy Donors. UMAP of post integration merged dataset of six samples after the nearest neighbors’ distance matrix construction for the experiments: A) Pre-filtered variable genes B) Permissive with post-merge selection of variable genes shown by experiment (left) and cluster (right) C) Permissive filtering only shown by experiment (left) and cluster (right). D) Dot plot of the most variable genes per cluster for permissive filtering. E) Top 5 informative genes per cluster for clusters in 5C.

When experiments pre-filtered for highly variable genes were used for the Harmony integration, the resulting merged dataset contained 68847 cells and only 218 genes, confirming the high heterogeneity of platelets. For permissive filtering, the merge yielded 16446 genes, and later applied filtering for variable genes resulted in 3000 genes. With pre-filtered data, the merge produced extreme clustering of cells, with more than 1700 well-integrated clusters presented using low-resolution Leiden, and did not produce any notable insights, that were additive to the finding of complex platelet heterogeneity (Figure 5A). Reasonable cluster numbers were observed within permissive filtering merge, 13 clusters with the selection of variable genes post permissive merge (Figure 5B), and 7 clusters without post selection for variable genes (Figure 5C). A dot plot and heatmap of the ranked differentiating genes in each of the clusters is shown in Figure 5 (D-E). Lastly, a dot plot of the ranked differentiating genes in each of the clusters in permissive filtering with later selection of variable genes, and a heat map of the ubiquitous genes identified earlier, and a list of well-established platelet-associated genes, representing a variety of functions including platelet activation, signaling, granule secretion, coagulation, cytoskeletal regulation, and megakaryocyte differentiation are presented in Supplementary Figure 3. The gene lists mentioned are presented in the Supplementary Table 1. CellTypist cell-type annotation, majority voting on clusters, generated for the integrated datasets which are presented in Supplementary Figure 4, indicating wide mischaracterization of platelets using modern cell-type annotation tools.

These findings further support highly heterogeneous gene expression of platelets, and the possibility that platelets may have been misidentified or overlooked in previously published studies which employed currently available pipelines for annotation, despite their presence. Furthermore, our findings suggest that prior platelet transcriptomic datasets are likely confounded due to the presence of other cell types.

## Discussion

We successfully conducted single-cell sequencing on platelets from seven healthy donors. In the initial group, we sequenced 50,000 acridine orange-high platelets, ensuring sufficient RNA content for analysis. Our results showed that the platelet transcripts closely mirrored those observed in bulk RNA sequencing. However, we were surprised by the significant heterogeneity in gene expression, with most platelets lacking common platelet-associated proteins, such as ITGA2B (CD41), GP1B (CD42b), and TREML1. To confirm that we were sequencing platelets, we employed antibody-based sequencing (AbSeq), which verified the low expression of these typically abundant transcripts. Further analysis revealed that platelet heterogeneity complicates cell annotation, with MSigDB algorithms misclassifying platelets as various other blood-associated cells, including podocytes and mesangial cells.

Compared to nucleated cells, platelets have greatly reduced amounts of RNA and a significantly higher percent of mitochondrial genes(22). Traditional guidelines for RNA sequencing used for somatic cells would eliminate many of the platelets whose number of transcripts and mitochondrial genes would exclude them from consideration following standard QC parameters. We have successfully used acridine orange to identify RNA containing platelets and sequenced the platelet transcriptome utilizing the BD Rhapsody™ platform (Figure 1). We found each platelet to have between 100 and 250 mean unique transcripts per cell. We proposed the use acridine orange to identify platelets that have the highest amount of RNA for sequencing and establish a platelet specific pipeline for researchers to follow; however, these platelets would be representative of only a small percentage of circulating cells. Using AbSeq with platelet specific surface markers, we achieved robust RNA sequencing of platelets suggesting acridine orange is not necessary to identify RNA containing platelets. Interestingly, there was a shift in the transcripts identified which is consistent with bulk sequencing of platelets sorted based on levels of thiazole orange (similar to acridine orange) staining(6). For example, the top transcript identified in the acridine orange high platelets of OU-HD1 and 2 was RORA. In the platelets from OU-HD3, whose acridine orange sorted platelets were not as high as HD1 or 2, RORA was not the most abundant transcript as was the case for the four AbSeq donors. While these results suggest that there are different populations of platelets based on RNA content, it also brings forth the question of if these differences have functional associations.

While previous studies have reported single-cell sequencing of platelets, our data suggest that these published algorithms may introduce significant bias. For example, studies on platelets from patients with periodontitis and diabetes used algorithms that selected transcripts based on platelet-specific proteins and required over 250 transcripts, often associating platelets with peripheral blood mononuclear cells (PBMCs). Furthermore, these platelets had been cryopreserved which has been reported to result in formation of procoagulant platelets which have an altered transcriptomic profile(23). Our data, derived solely from isolated platelets using single-cell RNA sequencing, indicate that not all platelets express top genes like α2b/β3a (CD41/CD61), and most contain fewer than 250 transcripts. In addition, platelets that associated with monocytes have higher levels of activation characterized by increased surface CD62P expression(24). Our findings, based on isolated platelets, suggest that not all platelets express key genes like ITGA2B/ITGB3 (CD41/CD61), which are commonly used proteins for platelet identification. Collectively, this suggests that previous studies may have misidentified platelets or only captured a subset of activated platelets due to overly strict selection criteria. In contrast, our study provides an accurate and comprehensive single-cell analysis of platelets, aligning well with bulk RNA sequencing data.

Megakaryocytes have been found to make platelets in the lung and they are believed to be responsible for up to 50% of the peripheral platelet population(3). It has been observed that lung megakaryocytes possess distinct transcriptional profiles compared to those found in bone marrow. Based on our results, we did not identify a strong correlation between published megakaryocyte and our platelet transcriptome.

The transcriptional heterogeneity of platelets may reflect either remnants of the megakaryocyte’s RNA or the horizontal transfer of RNA from other cells, as suggested by prior studies on platelet RNA exchange(25). Over the platelet’s lifespan, its transcriptome could potentially reflect the individual’s health status. This has exciting implications for liquid biopsies in cancer detection where platelets have been identified as biomarkers of disease severity. Alternatively, the RNA in platelets might simply represent what’s left from megakaryocyte transcription, akin to survivors on a life raft escaping a sinking ship. Our pathway analysis supports this, revealing correlations with protein production machinery present in platelets. These findings open new avenues for understanding platelet biology and underscore the need for tailored approaches in platelet transcriptomics.

The findings presented in our study provide a foundation for understanding of heterogeneity of platelets. We highlighted the importance of careful planning of methodologic approaches for platelet transcriptomic investigations. Furthermore, we highlight limitations of current annotation pipelines for accurate identification of platelets. This misclassification may introduce multiple detractions to existing and prospective studies and lead to false conclusions. Additional experimental efforts with larger sample sizes are needed to further investigate the heterogeneity of platelet gene expression.

## Acknowledgments

This work was funded in part through National Institutes of Health grants from National Heart, Lung, and Blood Institute R01HL903303 (V. W.) K23HL151872 (D.R.L.), Harold Amos Medical Faculty Development Award/American Heart Association/Robert Wood Johnson (D. R. L.), CCTST Mentored Translational Grant (D.R.L.), University of Cincinnati DOIM Pilot Grant (D.R.L.), Startup Funds Oakland University (A.V. W.), and The Leona M. and Harry B. Helmsley Charitable Trust (T.O).

## Authorship and Conflict of Interest Statements

Conduct of experiments, CD, CF, AVW, DL; Writing of the manuscript WW, DL, AVW; Analysis of data, WW, TO, DL, AVW; Direction of studies WW, TO, DL, AVW.

There are not conflict of interest for WW, CD, CF, TO, AVW, or, DL.

