## Supplemental data for "The First Comprehensive Description of the Platelet Single Cell Transcriptome"

### Slide 1
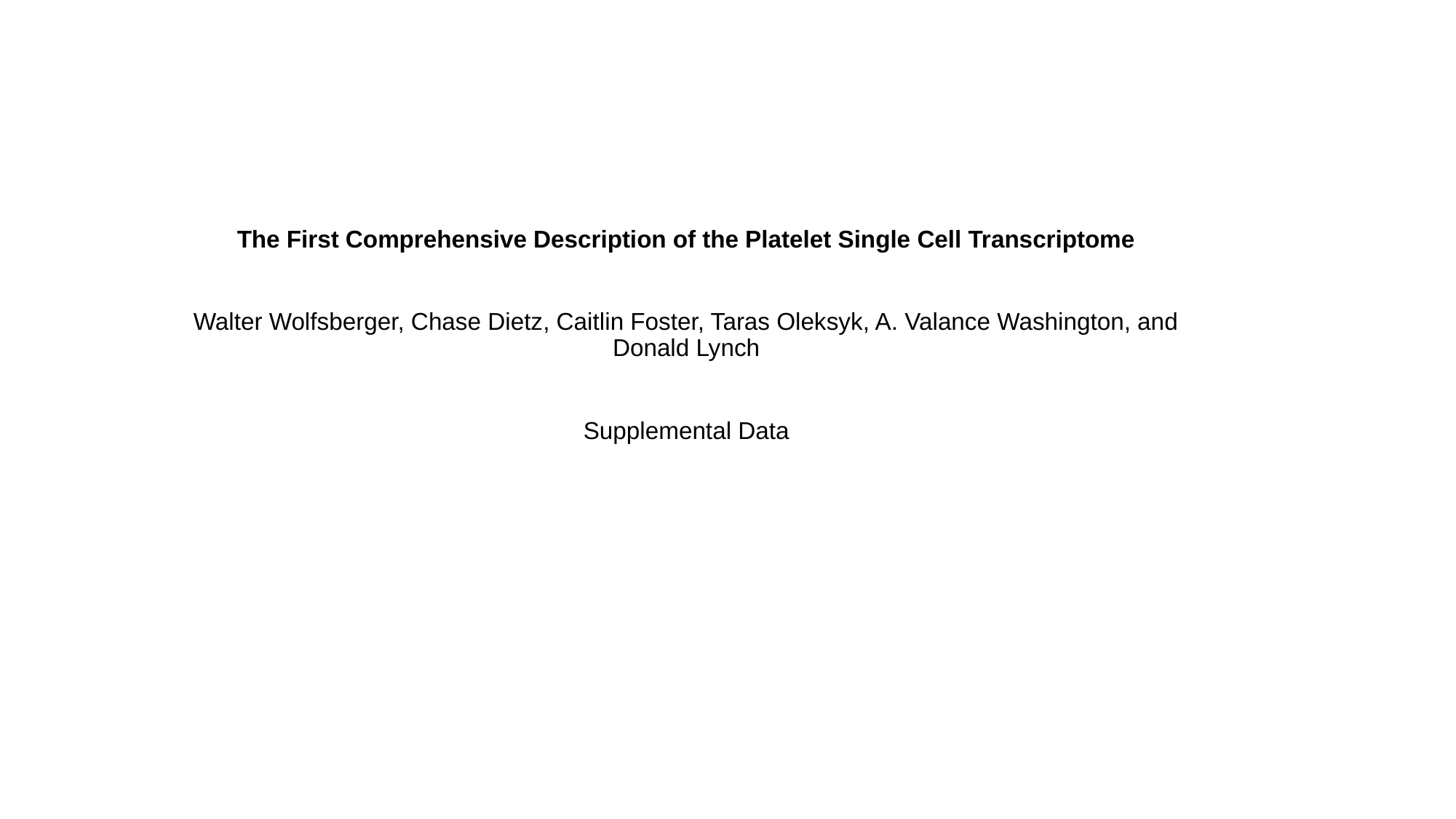

The First Comprehensive Description of the Platelet Single Cell Transcriptome
Walter Wolfsberger, Chase Dietz, Caitlin Foster, Taras Oleksyk, A. Valance Washington, and Donald Lynch
Supplemental Data

### Slide 2
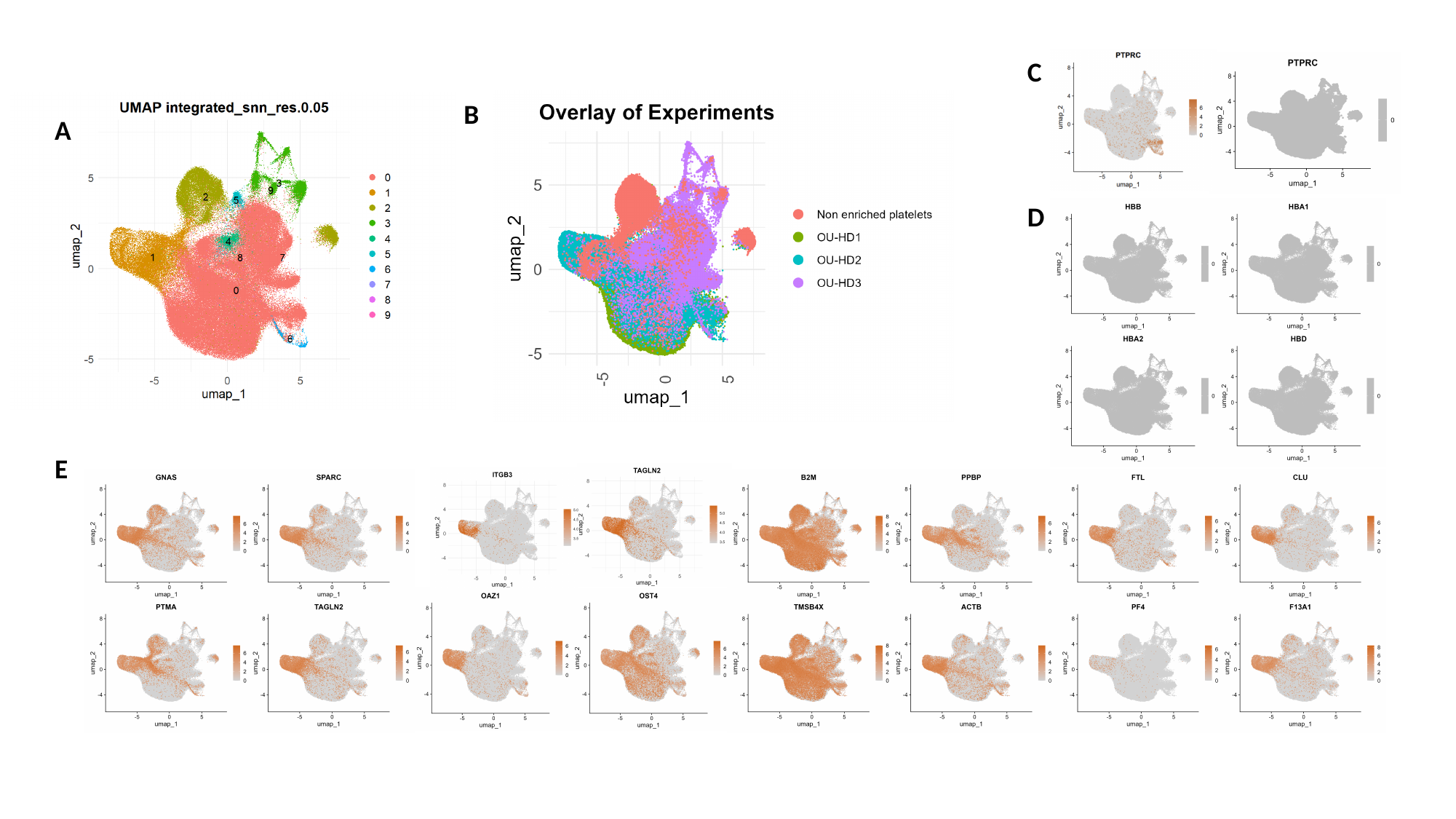

C
B
A
D
E

### Slide 3
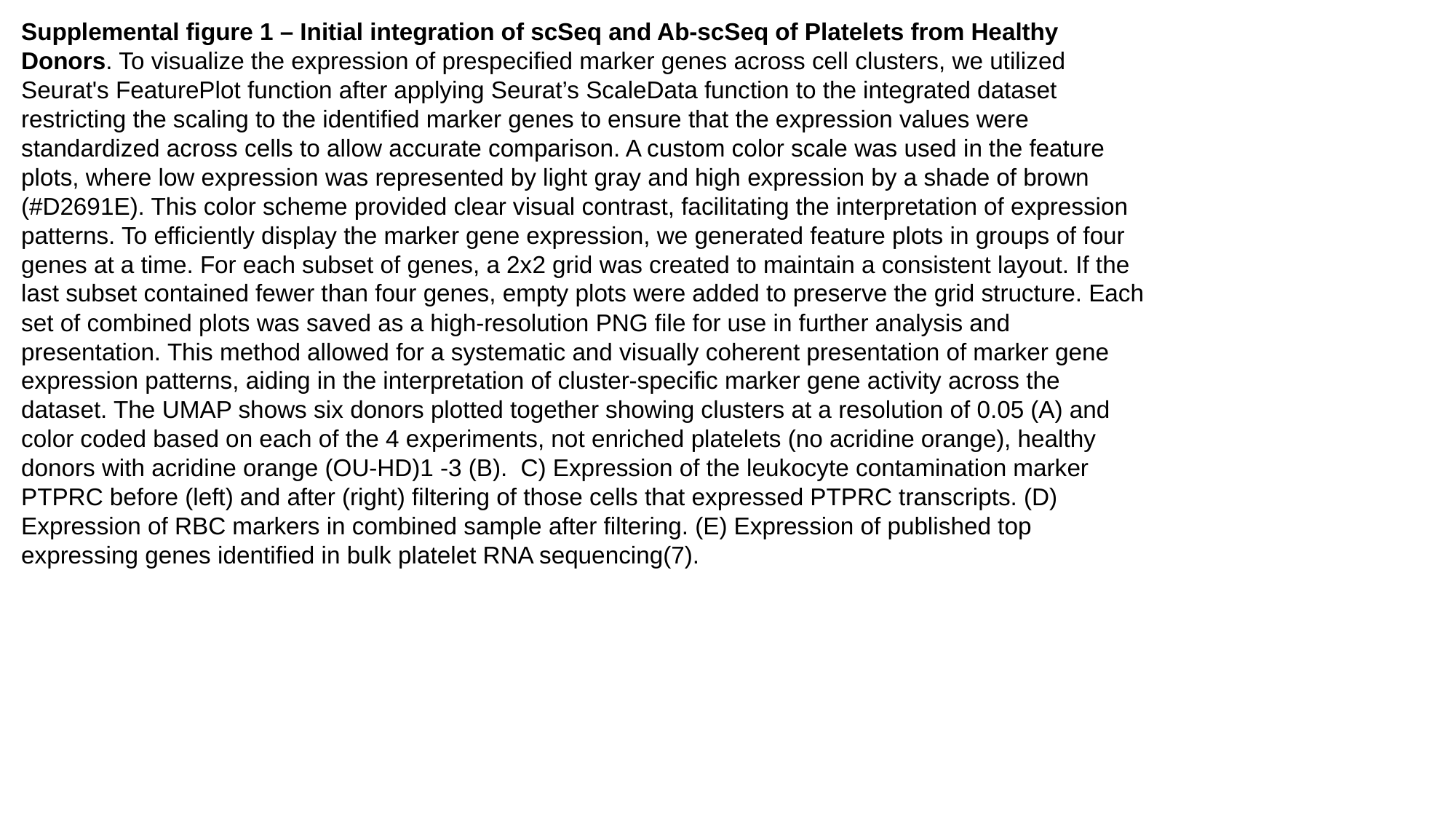

Supplemental figure 1 – Initial integration of scSeq and Ab-scSeq of Platelets from Healthy Donors. To visualize the expression of prespecified marker genes across cell clusters, we utilized Seurat's FeaturePlot function after applying Seurat’s ScaleData function to the integrated dataset restricting the scaling to the identified marker genes to ensure that the expression values were standardized across cells to allow accurate comparison. A custom color scale was used in the feature plots, where low expression was represented by light gray and high expression by a shade of brown (#D2691E). This color scheme provided clear visual contrast, facilitating the interpretation of expression patterns. To efficiently display the marker gene expression, we generated feature plots in groups of four genes at a time. For each subset of genes, a 2x2 grid was created to maintain a consistent layout. If the last subset contained fewer than four genes, empty plots were added to preserve the grid structure. Each set of combined plots was saved as a high-resolution PNG file for use in further analysis and presentation. This method allowed for a systematic and visually coherent presentation of marker gene expression patterns, aiding in the interpretation of cluster-specific marker gene activity across the dataset. The UMAP shows six donors plotted together showing clusters at a resolution of 0.05 (A) and color coded based on each of the 4 experiments, not enriched platelets (no acridine orange), healthy donors with acridine orange (OU-HD)1 -3 (B). C) Expression of the leukocyte contamination marker PTPRC before (left) and after (right) filtering of those cells that expressed PTPRC transcripts. (D) Expression of RBC markers in combined sample after filtering. (E) Expression of published top expressing genes identified in bulk platelet RNA sequencing(7).

### Slide 4
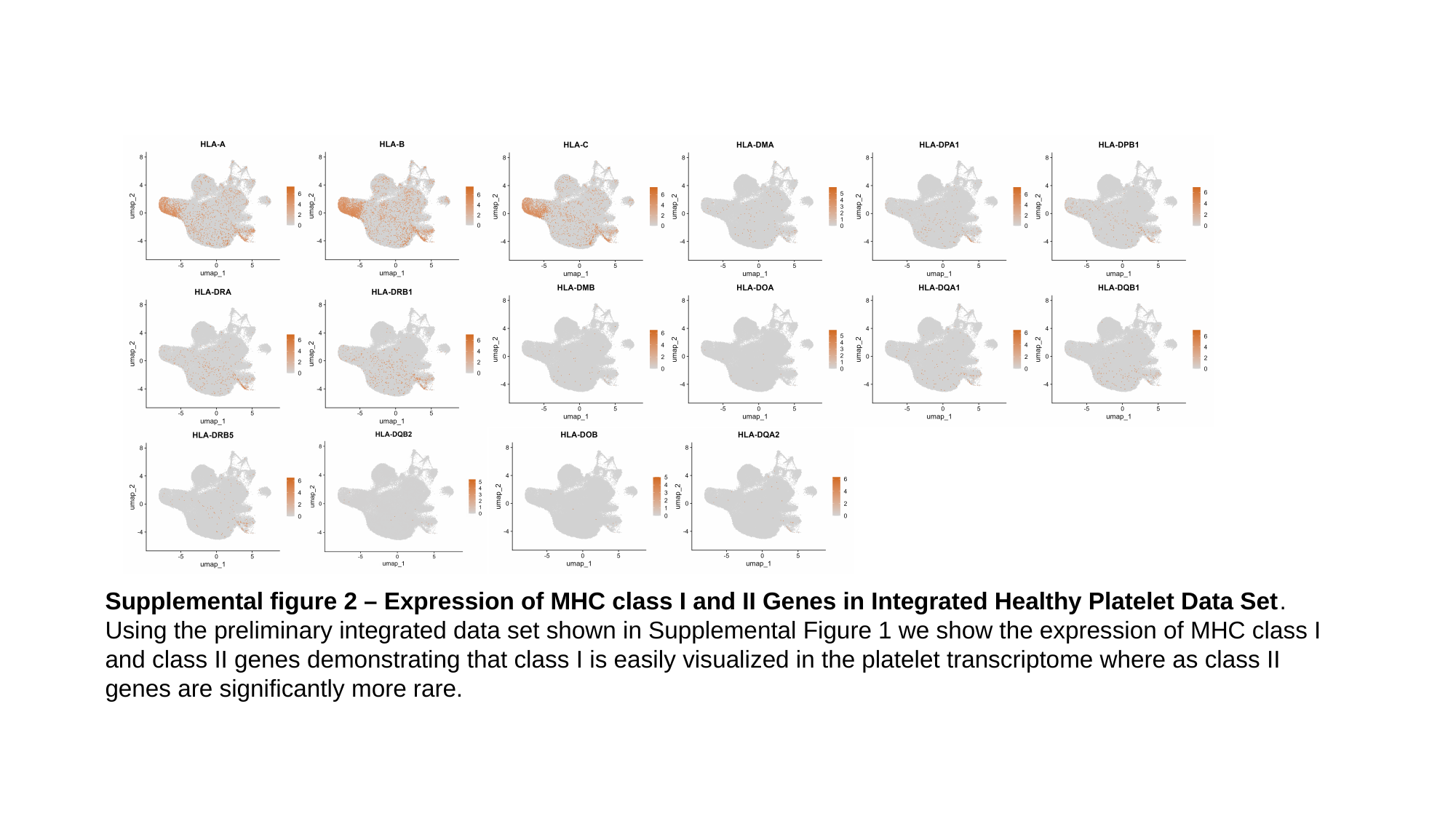

Supplemental figure 2 – Expression of MHC class I and II Genes in Integrated Healthy Platelet Data Set. Using the preliminary integrated data set shown in Supplemental Figure 1 we show the expression of MHC class I and class II genes demonstrating that class I is easily visualized in the platelet transcriptome where as class II genes are significantly more rare.

### Slide 5
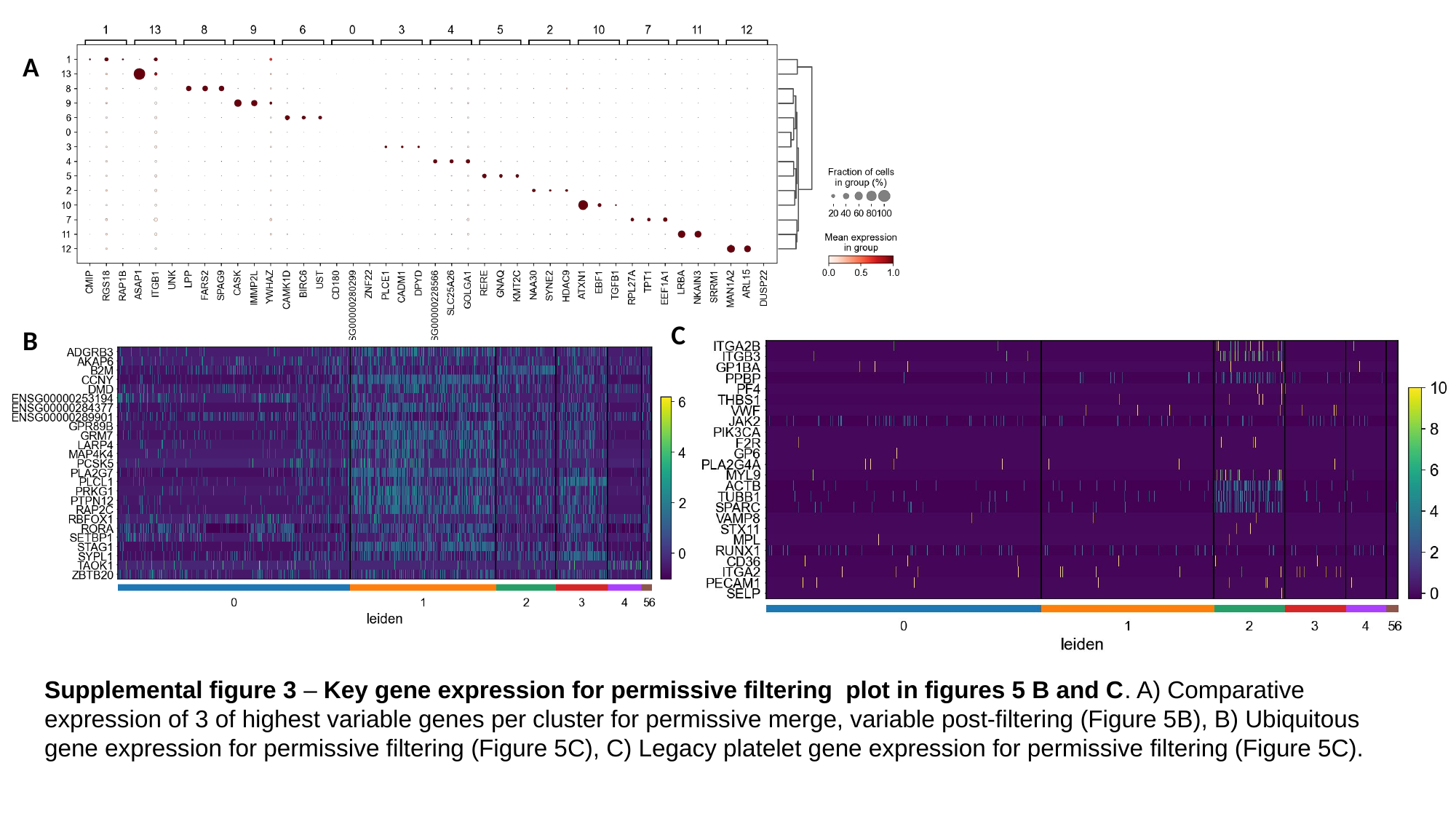

A
C
B
Supplemental figure 3 – Key gene expression for permissive filtering plot in figures 5 B and C. A) Comparative expression of 3 of highest variable genes per cluster for permissive merge, variable post-filtering (Figure 5B), B) Ubiquitous gene expression for permissive filtering (Figure 5C), C) Legacy platelet gene expression for permissive filtering (Figure 5C).

### Slide 6
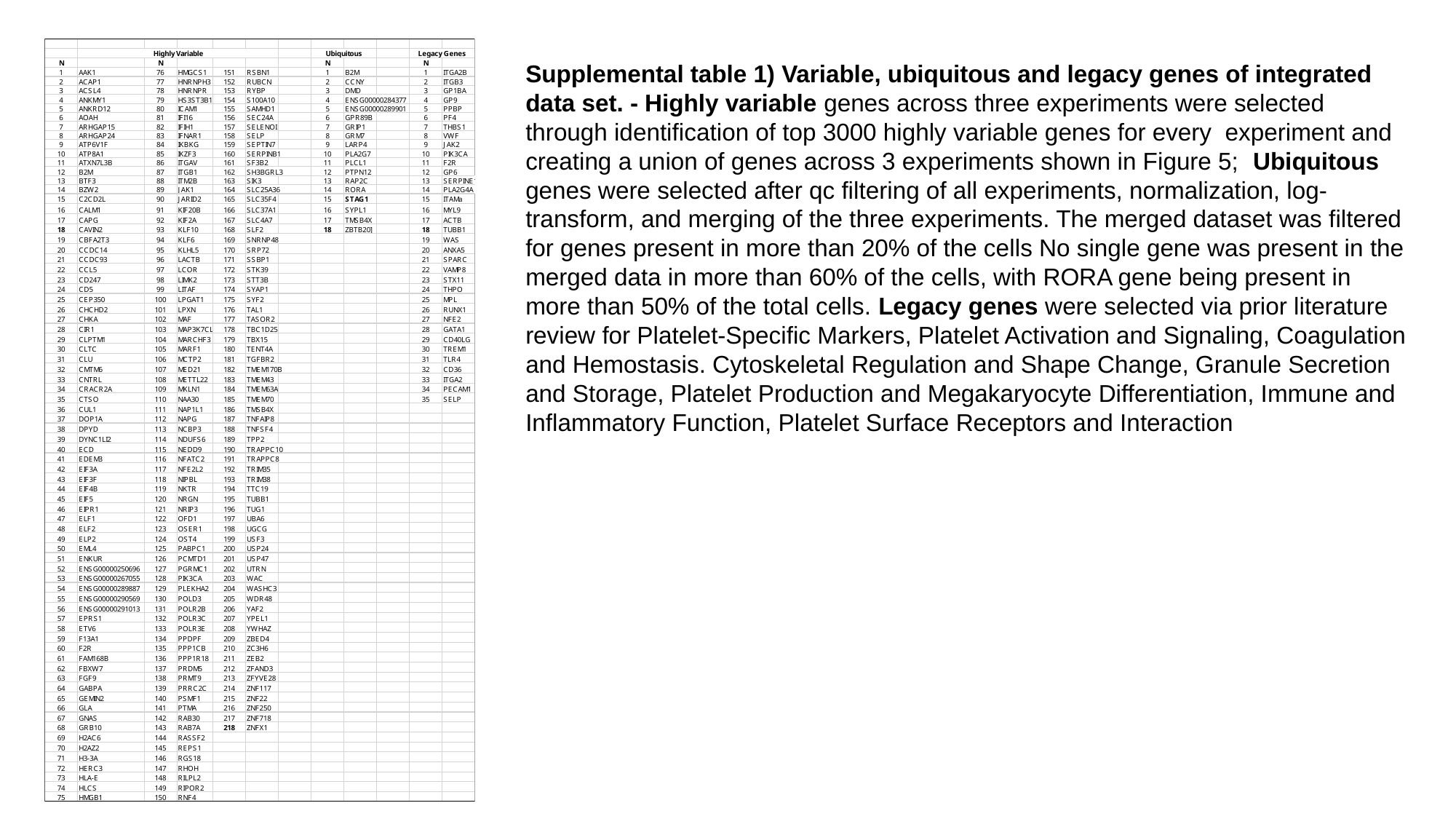

Supplemental table 1) Variable, ubiquitous and legacy genes of integrated data set. - Highly variable genes across three experiments were selected through identification of top 3000 highly variable genes for every experiment and creating a union of genes across 3 experiments shown in Figure 5; Ubiquitous genes were selected after qc filtering of all experiments, normalization, log-transform, and merging of the three experiments. The merged dataset was filtered for genes present in more than 20% of the cells No single gene was present in the merged data in more than 60% of the cells, with RORA gene being present in more than 50% of the total cells. Legacy genes were selected via prior literature review for Platelet-Specific Markers, Platelet Activation and Signaling, Coagulation and Hemostasis. Cytoskeletal Regulation and Shape Change, Granule Secretion and Storage, Platelet Production and Megakaryocyte Differentiation, Immune and Inflammatory Function, Platelet Surface Receptors and Interaction

### Slide 7
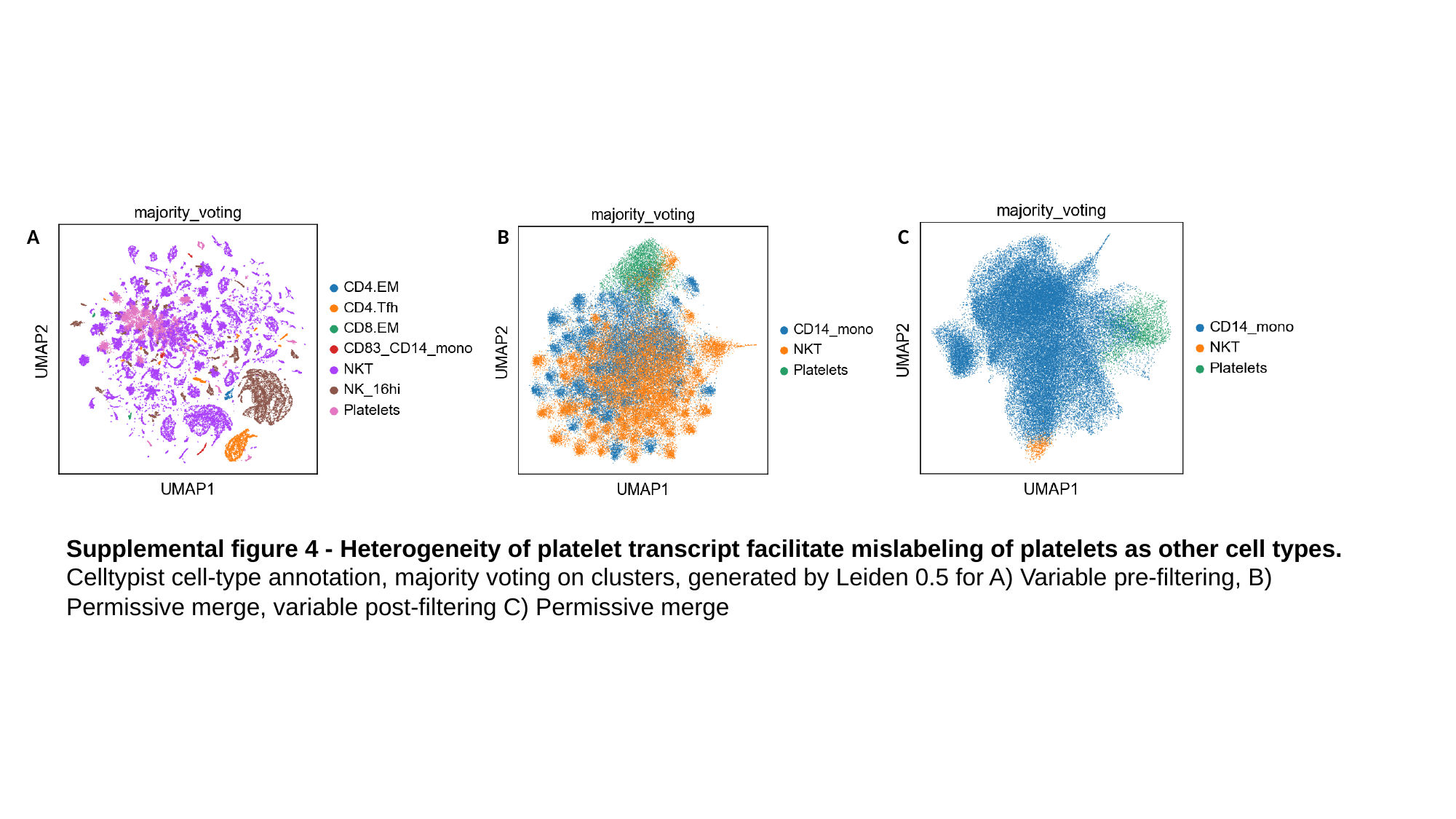

C
A
B
Supplemental figure 4 - Heterogeneity of platelet transcript facilitate mislabeling of platelets as other cell types. Celltypist cell-type annotation, majority voting on clusters, generated by Leiden 0.5 for A) Variable pre-filtering, B) Permissive merge, variable post-filtering C) Permissive merge
